# Single-hit genome edition for expression of single-chain immunoglobulins by edited B cells

**DOI:** 10.1101/2022.06.02.494471

**Authors:** Natsuko Ueda, Marine Cahen, Christophe Sirac, Anne Galy, Jérôme Moreaux, Yannic Danger, Michel Cogné

**Affiliations:** INSERM U 1236, University of Rennes 1, Etablissement Français du Sang, 35000 Rennes, France; INSERM U1262, CNRS UMR 7276, Limoges University, Control of the B-cell Response & Lymphoproliferation, 87025 Limoges, France; Université Paris-Saclay, Univ Evry, Inserm, Genethon, Integrare research unit UMR_S951, 91000, Evry, France; CNRS-UM UMR 9002, Institute of Human Genetics, 34090 Montpellier, France

## Abstract

Lymphocytes have become attractive agents for adoptive immunotherapy but only the reformatting of T cells is efficiently mastered. Despite some recent breakthroughs, B cells remain challenging targets, with regard to both their long-term survival after *in vitro* manipulation and the rewiring of immunoglobulin (Ig) expression. Working on these two aspects, we have designed a new format of single-chain Ig (“scFull-Ig”) coding cassette, the insertion of which at a single genomic position can redirect B cells toward a new antigen specificity, while preserving all functional domains of the B cell receptor. Precise genomic edition at a single locus then provides the most efficient and safe strategy to both disrupt endogenous Ig expression while encoding a new Ig paratope. As proofs of concept, functionality of such scFull BCR was validated by checking specific binding of two different classical targets of tumor immunotherapy, HER2 and CD20. Once the strategy validated in cell lines, it was also validated in primary B cells, again showing successful engineering of BCR expression. These results contribute to pave the way for future B cell-based adoptive cell therapy.

## INTRODUCTION

Efficient monoclonal antibodies (mAb) are available for the treatment of multiple chronic diseases and notably cancers. The high specificity of such biotherapies hereby limits their side effects in comparison to conventional treatments such as chemotherapy of cancer. However, mAbs can in some cases yield poor response or resistance, especially in those patients undergoing the lowest serum trough concentration (C_min_)^1^. Repeated and frequent injections of mAb are then required to prevent relapses, which can request prolonged and costly care and be restrictive for these patients. This is especially true for mAbs with rapid serum clearance, such as panitumumab or cetuximab or bispecific antibodies^1^. Strategies that could result into permanent infusion of such antibodies, for instance as endogenously produced by the patient’s cells, would thus represent a strong asset. The usage of some antibodies of high therapeutic interest but with chemical instability or manufacturing contingencies could also be rescued by direct *in vivo* production in the patient’s body. This strategy could also be pertinent for conferring long-term immunity to some immunodeficient patients with a preserved B cell pool but poorly responsive to classical vaccination.

B lineage cells are indeed the best-suited cell type for the correct folding and secretion of immunoglobulins (Ig) under their various membrane bound or secreted formats^2^. While plasma cells are able to secrete large amounts of soluble antibodies, memory B cells can in addition establish an immune memory and be responsive to antigen (Ag) re-challenge, hereby contributing to protection against recurrent infections. Editing Ig expression in such cells could thus potentially endow patients with long-term immune response to a specific Ag. Several recent studies have shown that CRISPR- edition of the B cell genome is feasible in primary B cells and could reformat Ig production on purpose. Regarding cancer immunotherapy and learning from the success of CAR-T cells, which have now permitted complete remission in many patients and demonstrated their ability to safely integrate the patients’ immune system for years^3–5^, the conditions that could allow successful adoptive cell therapy based on B cells are thus worth being explored and optimized. In such a strategy, B cells taken from an individual would be genetically modified to produce Ig of a desired specificity and then injected back to the original donor. Successful engraftment of such engineered cells would then be expected to yield long-term stable infusion of a therapeutic mAb into body fluids, and/or an immune memory artificially built against a tumor Ag.

We used for this purpose CRISPR-mediated gene knock-in (KI), as the best tool for precise gene edition^6,7^, in order to hijack the innate capacities of B cells to produce Ig molecules. The insertion site for the KI was chosen within the Ig heavy chain (IgH) locus in order to obtain simultaneously both the disruption of the endogenous IgH synthesis and the integration of the KI cassette in a position mimicking all the normal features of Ig expression. These include B-lineage-specific transcription, upregulated transcription in activated B cells and plasma cells, regulation of alternative splicing between membrane-type and secreted-type transcripts, ability to undergo class switch recombination upon recombination between the endogenous Sμ region and a downstream S region from the IgH locus.

*In vitro* cell culture and maintenance of B cells together with genetic engineering remain challenging. We optimized B cell culture conditions and CRISPR edition. We now describe an efficient strategy to gene edit human primary B cells in order to produce a complete Ig of a given specificity after a single genetic edition. The genetic engineering of Ig theoretically requires the modification of two loci, for the heavy (H) and light (L) chains. To express the desired Ig and disrupt endogenous Ig production after a single gene modification, we designed a new single chain format of a whole Ig, the “scFull-Ig”. This format ensures stoichiometric expression of both Ig chains and precludes any mispairing with an endogenous Ig light chain. It also carries the advantage of positioning both the V_H_DJ_H_ region and V_L_J_L_ region immediately downstream of the V_H_ promoter, at a position accessible to somatic hypermutation (SHM), while constant regions lie downstream. With this strategy, modified primary B cells proved to be able to successfully express scFull-Ig format BCR membrane molecules, hereby acquiring the relevant Ag-binding activity.

## MATERIALS AND METHODS

### scFull-Ig structure validation

At the protein level, we wished to produce full-length Ig under a single chain format by using two linker (Lnk) sequences for assembling Ig domains in the following order: [VH-Lnk-VLQ-Lnk-CH]. For building adequate gene cassettes encoding such an scFull-Ig, we thus assembled the coding sequences from a chosen mAb in the following 5’ to 3’ order: V_H_ leader exon, V_H_DJ_H_ exon, a first linker (GGTGGTGGTGGTTCTGGTGGTGGTGGTTCTGGCGGCGGCGGCTCCAGTGGTGGTGGATCC), V_L_J_L_exon, C_L_ exon, followed by a 2^nd^ linker (TCTGGTGGCGGTGGCTCGGGCGGAGGTGGGTCGGGTGGCGGCGGATCA). This Ag-specific upstream part of the gene construct was either followed with a C_H_ coding sequence in order to directly encode a complete IgG, or flanked with a splice donor site and its 3’ flanking sequence identical to that from a J_H_ element, in order to be expressed after a KI upstream of an endogenous C_H_ gene. Adequately designed sequences were synthetized (Genecust) and cloned in a pcDNA3 expression vector. HEK293T cells were transfected with different constructs to validate expression. After 72h, the supernatants were collected, cleared by centrifugation, and stored at 2-8°C for further experiments. Supernatants were analyzed by western blot after reduction by 2-mercaptoethanol (Bio-Rad). Electrophoresis was carried out using nUView Tris-Glycine 8-16% gels (NuSep) and proteins were then transferred on a PVDF membrane. Membranes were stained using anti-κ (Southern biotech) and anti- λ antibodies (Southern biotech) followed by an HRP-conjugated donkey anti-goat IgG antibody (Santa Cruz Biotechnology), or using HRP-conjugated goat anti-human IgG antibody (Sigma). Immunoblots were revealed using SuperSignal West Pico PLUS Chemiluminescent Substrate (Thermo Scientific). Controls used were Rituximab (MabThera, Roche) and Pertuzumab (Perjeta, Roche).

ADCC assays were carried out using CD16-expressing reporter cells from the Jurkat-Lucia NFAT-CD16 cell line (jktl-nfat-cd16, Invivogen) following manufacturer’s protocol. Briefly, target cells (MCF7 or hCD20^+^ EL4) were incubated with antibodies for one hour, after which Jurkat-Lucia NFAT-CD16 cells were added. After incubating for 6 hours, the supernatant was transferred into a white microplate, and QUANTI-Luc solution (Invivogen) was added. The light signal reporting Jurkat cell activation was quantified using a luminometer.

### Cell culture

BL41 cells were grown in RPMI 1640 GlutaMAX medium (Gibco) supplemented with 10% FBS (Gibco), 1 mM Sodium Pyruvate (Gibco) and Penicillin-Streptomycin-Glutamine (1 unit/ml penicillin, 1 μg/ml streptomycin, 2.92 μg/ml L-glutamine, Gibco). One day before CRISPR experiments, cells were seeded at 0.2.10^6^ cells/ml to be in an exponential phase of growth.

HEK293T cells were grown in DMEM medium (Gibco) supplemented with 10% FBS (Capricorn) and PS- G. Cells were seeded 2 or 3 days before transfection experiment by MACSfectin reagent (Miltenyi Biotec), to achieve cell density recommended by the supplier.

### Primary B cell isolation and culture

Buffy coats from healthy volunteers were obtained from the Etablissement Français du Sang (Rennes, France) (after informed consent, and under agreement #AC-2019-3853). B cells were isolated using either the StraightFrom™ Buffy Coat CD19 Microbead Kit or the Human B cell Isolation Kit II (both from Miltenyi Biotec), following the manufacturer’s protocols. When using Human B cell Isolation Kit II, peripheral blood mononuclear cells (PBMCs) were isolated using Histopaque (Sigma) or Lymphocyte Separation Medium (Eurobio Scientific) and red blood cells were lysed with NaCl solution during washing steps. B cells were then cultured in RPMI 1640 GlutaMAX medium (Gibco) supplemented with 10% FBS (Gibco), 1 mM Sodium Pyruvate (Gibco) and PS-G (Gibco). During four days, B cells were stimulated with 1 μg/ml CpG oligodeoxyribonucleotide 2006 (Miltenyi Biotec), 2.4 μg/ml F(ab’)_2_ Fragment Goat Anti-Human IgA + IgG + IgM (H+L) (Jackson ImmunoResearch), 50 U/ml recombinant IL-2 (R&D Systems) and 100 ng/ml recombinant human soluble CD40L (Immunex) (“1^st^ step stimulation cocktail”) at 0.75.10^6^ cells/ml. At day two, IL-10 (R&D Systems) was added at 5 ng/ml. At day four, activated B cells were washed and cultured at 0.5.10^6^ cells/ml with 50 U/ml IL-2 (R&D Systems), 5 ng/ml IL-4 (R&D Systems) and 12 ng/ml IL-10 (R&D Systems) in order to promote the differentiation into plasmablasts (“2^nd^ step differentiation cocktail”)^8^. To obtain plasma cells, cells were washed at day 7 and cultured at 0.5.10^6^ cells/ml with 50 U/ml IL-2 (R&D Systems), 250 U/ml IFN-α (R&D Systems), 20 ng/ml IL-6 (Peprotech) and 12 ng/ml IL-10 (R&D Systems).

### Donor DNA preparation and gene edition

Plasmids containing templates for CRISPR-mediated KI were assembled using the NEBuilder HIFI assembly kit (New England Biolabs) starting from PCR amplified fragments. Donor DNA for CRISPR-mediated KI were prepared from these plasmids by PCR amplification. PCR products were then purified using NucleoSpin Gel and PCR Clean-up Kit (Macherey-Nagel), with a final target concentration of 500-1000 ng/μl.

After comparing several sites in-between JH6 and the Eμ enhancer of the IgH locus, we focused our experiments by targeting the following sequence, GGAAAGAGAACTGTCGGAGT, with an appropriate sgRNA. For CRISPR edition, 100 pmol of Cas9 (IDT) were incubated with 500 pmol of sgRNA (Synthego) for 15 min at RT. Cells were nucleofected using Cell Line Nucleofector Kit V (Lonza) for cell lines and P3 Primary Cell 4D-Nucleofector X Kit L (Lonza) for primary B cells. 1.10^6^-3.10^6^ cells were centrifuged 10 min at 90 g and resuspended in 100 μl supplemented Nucleofector solution. The Cas9/sgRNA mix was then added to the cell suspension and cells were transfected with Nucleofector I program X-01 for cell lines and Nucleofector 4D program EO-117 for primary B cells. 900 μl culture medium was added immediately after transfection and cells were left resting for 5-10 min. Cells were then transferred into culture plates in 5 ml medium for cell lines or 1.5 ml for primary B cells.

When KI was performed, the donor DNA was added to the Cas9/sgRNA mix before addition to cells. For primary B cell transfections, if required, PGA (Sigma) was added to sgRNA before mixing with Cas9 at a final concentration of 4 mg/ml, Electroporation Enhancer (IDT) was added to the Cas9/sgRNA mix at a final concentration of 4 μM just before addition of cells, and HDR Enhancer (IDT) was added to the culture medium at a final concentration of 30 μM after transfection and removed by medium renewal after 40 h.

### DNA and RNA analysis

Correct CRISPR/Cas9 targeting of edited cells was checked by analyzing DNA through Tracking of Indels by Decomposition (TIDE) analysis and by sequencing edited transcripts. Cell pellets were lysed using QuickExtract DNA Extraction Solution (Lucigen) following supplier’s protocol. The region of interest was amplified by PCR, and Sanger Sequencing was performed on the product by Genewiz. Returned sequences were analyzed by TIDE (http://shinyapps.datacurators.nl/tide/).

RNA was extracted from transfected cells using NucleoSpin RNA Plus XS kit (Macherey-Nagel). cDNA of the IgH transcripts was synthesized and amplified by PCR, using one primer binding specifically to the scFull-Ig KI cassette and the other one to either μ or α constant domain. PCR products were then sequenced (Genewiz) to confirm the correct splicing of the inserted sequence on the endogenous CH1 constant domain of the IgH locus.

All primers used in the study are listed in Supplementary information.

### Flow cytometry

Cell surface staining for the anti-HER2 BCR was performed with Recombinant Human ErbB2/Her2 Fc Chimera Protein (R&D Systems) and APC anti-His Tag antibody (Biolegend) for 30 min each. DAPI (Serva) was used to mark dead cells. Stained cells were analyzed on a CytoFLEX flow cytometer (Beckman Coulter).

### Statistical analysis

Statistical analyses were performed using Mann-Whitney tests with GraphPad Prism 8 software. Data shown as mean ± SD, * p<0.05, ** p<0.01.

## RESULTS

### The scFull-Ig design allows efficient secretion

We designed and validated a new single-chain format of Ig, the scFull-Ig (Figure 1.A). This design connects via a linker the variable domain of the heavy chain with the variable and constant domains of the _k_ light chain, then followed by a second linker and the full constant region of the Ig heavy chain. In order to first validate the functionality of this antibody format, we produced the secreted forms of scFull mAb targeting CD20 or HER2. Recombinant scFull-IgG was produced by transfecting HEK293T cells with the adequate expression vector, which included a cassette successively linking a V_H_ promoter, the rituximab or pertuzumab V_H_DJ_H_ exon, a first linker, the complete κ light chain, a second linker ([VDJ_ritux_/_pertu_-Lnk-VκJκ_ritux_/_pertu_-Cκ-Lnk]) and finally the complete coding sequence of a human g1 heavy chain constant domains. We successfully produced scFull-IgG secreted in supernatants with concentrations reaching 374.1 ng/ml for anti-CD20 scFull-IgG and 364.4 ng/ml for anti-HER2 scFull-IgG. Supernatants containing scFull mAb were purified on protein A column for further characterization, with the same protocol as classical human IgG. Expression of Ig of the expected structure was confirmed by western blot using anti-hIgG, anti-κ and anti-λ antibodies (Figure 1.B). All stainings confirmed a single-chain structure linking heavy and light chains. We assessed and validated the binding of the anti-CD20 scFull-Ig on hCD20^+^ EL4 target cells (Figure 1.C).The functionality of these scFull-Ig was tested by assessing their capacity to induce antibody-dependent cellular cytotoxicity (ADCC). As for rituximab and pertuzumab, anti-CD20 scFull-IgG and the the anti-HER2 scFull-IgG were able to specifically induce ADCC against cells carrying the target antigen (*i.e*. hCD20^+^ EL4 cells, and MCF7 cells, respectively) even when used at low doses in the range of 10 ng/ml (Figure 1.D). Taken together, these data validate the functionality of the scFull-Ig format for producing Ag-specific antibodies.

**Figure 1.**
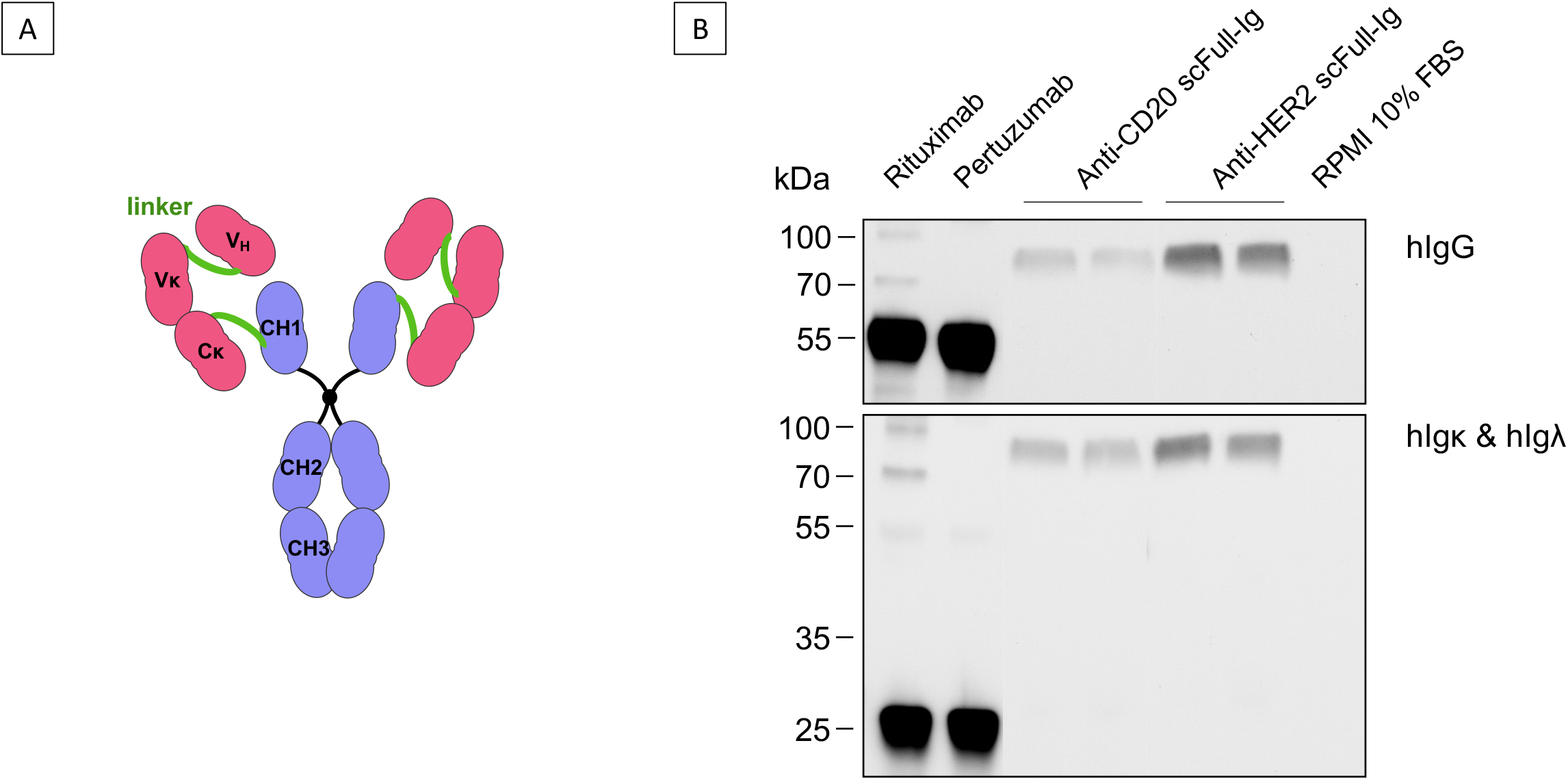
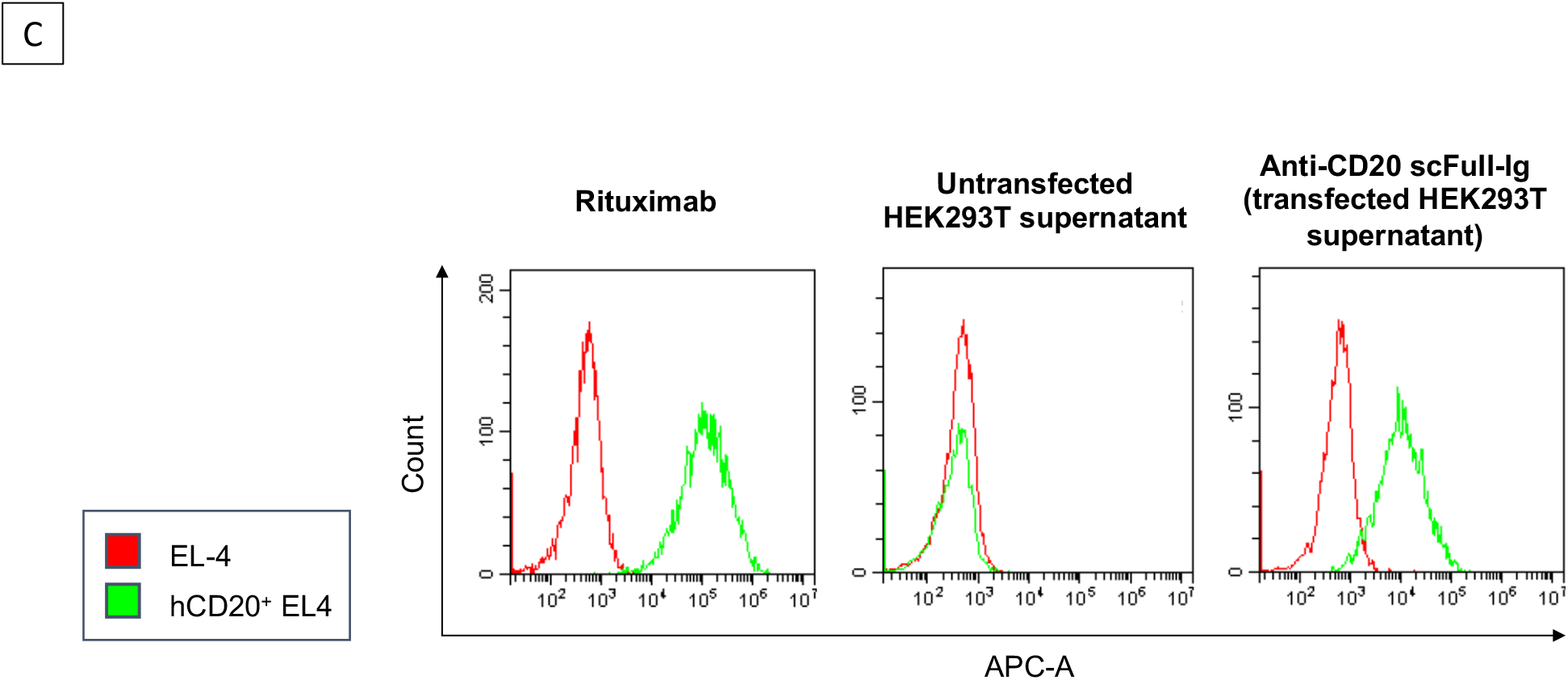
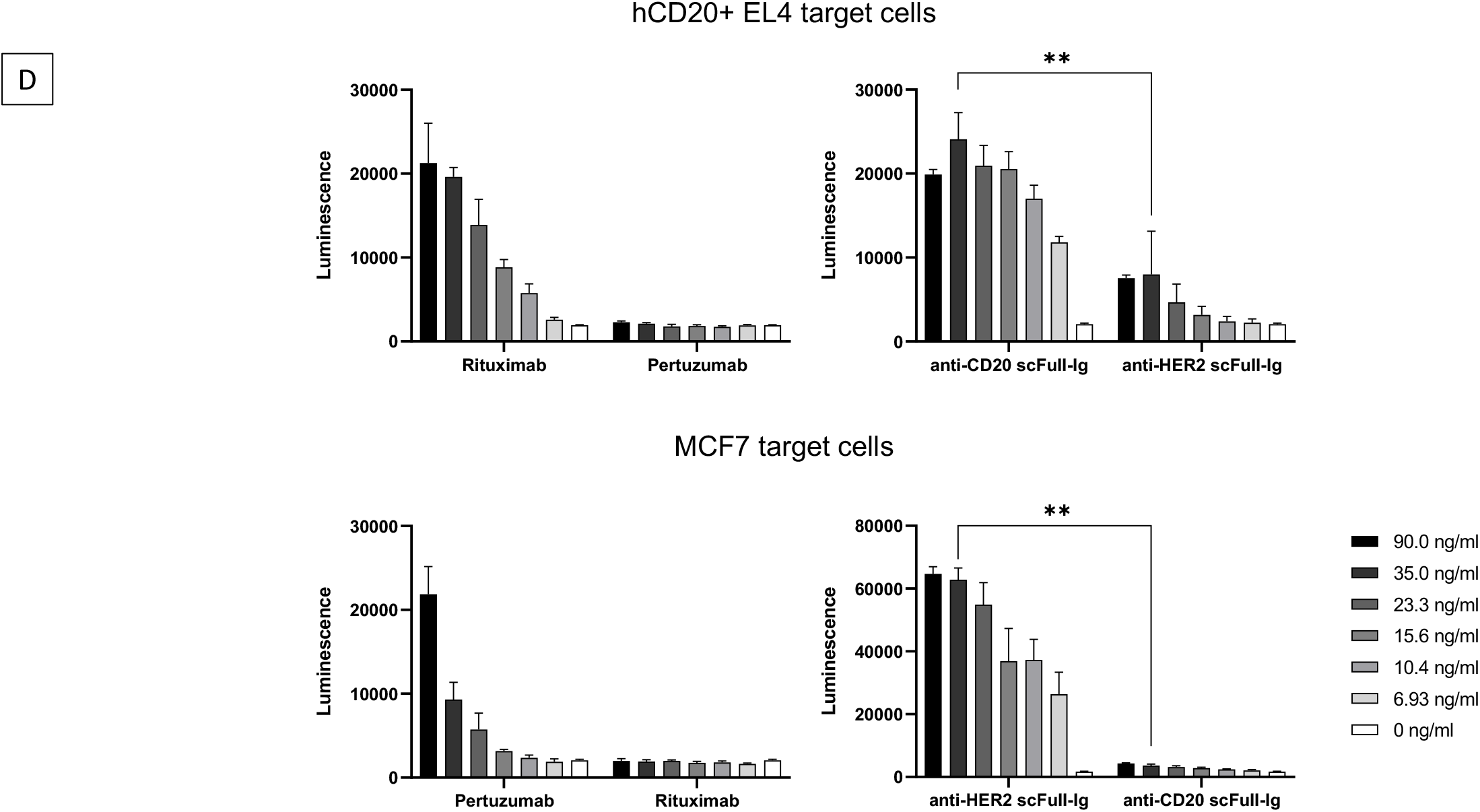
scFull-Ig. <u>(A)</u> Structure of an scFull-IgG. The scFull-Ig design combines all the domains from a complete Ig but under a single-chain format. The IgH variable domain is connected via a linker (*green*) to the light chain, itself followed by a second linker and the full IgH constant part. For CRISPR-mediated expression in B cells, the KI cassette only encodes for the IgH variable domain and the IgL sequences (*pink*). It is followed by a splice donor site to allow splicing on the endogenous genes encoding for the IgH constant exons (*blue*). <u>(B)</u> Detection of scFull-IgG expression by western blot using anti-hIgG, anti-hIgκ and anti-hIgλ antibodies confirms expression of a single-chain molecule of the expected size. <u>(C)</u> Validation of target binding by anti-CD20 scFull-IgG secreted by transfected cells. hCD20^+^ EL4 cells were used as targets, and using WT EL4 cells as a control, staining was revealed with a fluorescent anti- IgG secondary antibody. <u>(D)</u> Validation of functionality (ADCC). scFull-IgG secreted by transfected cells induced ADCC on target cells.

### Knock-in of a single scFull-Ig cassette allows efficient BCR expression in a B cell line

Besides the advantages of a single-chain for ensuring stoichiometry and physical association of the paired H/L structure, the scFull format was designed in order to allow recombinant Ig production through a single CRISPR-mediated genomic edition, at the IgH locus. The [VDJ-Lnk-VκJκ-Cκ-Lnk] cassettes, preceded by a VH promoter, were introduced in-between the J_H_ region and the Eμ enhancer (Figure 2.A). Insertion at this location allows the expression of the edited Ig while concomitantly disrupting expression of the endogenous Ig. We preliminarily validated the high accessibility of this genomic location for CRISPR edition in the BL41 lymphoma cell line, using an HDR template which included a pVH-promoter-tdTomato reporter cassette. This experiment also validated the efficacy of 500 bp-long homology arms which were finally designed with terminal truncated Cas9 target sequences (tCTS) at both ends of the HDR template to optimize HDR^9^. This altogether regularly yielded fluorescence with about 30% of fluorescent cells (Sup Figure 1). Based on this result and using the same HDR flanking arms, we then moved on to the KI of scFull-Ig expressing cassettes at the very same genomic location. In order to be appropriately spliced onto endogenous IgH constant genes, the scFull- Ig gene cassettes [VDJ-Lnk-VκJκ-Cκ-Lnk] previously validated above, were then also flanked by a downstream J_H_ splice donor site (J_H_3’splice). In such a configuration, the [VDJ-Lnk-VκJκ-Cκ-Lnk] can be spliced on the immediate downstream constant gene and replace the endogenousV_H_DJ_H_ exon in IgH transcripts (Figure 2.A). A KI at this position should affect neither the process of alternate splicing of membrane-type *vs* secreted type IgH C transcripts, nor the process of class switch recombination (CSR) of the locus. In cells undergoing CSR and/or plasma cell differentiation, such a KI is thus expected to yield (as for a normal endogenous IgH gene), multiple clonally related BCR and secreted Ig formats, related to the various Ig classes but sharing the same clonotypic Ag specificity.

**Figure 2.**
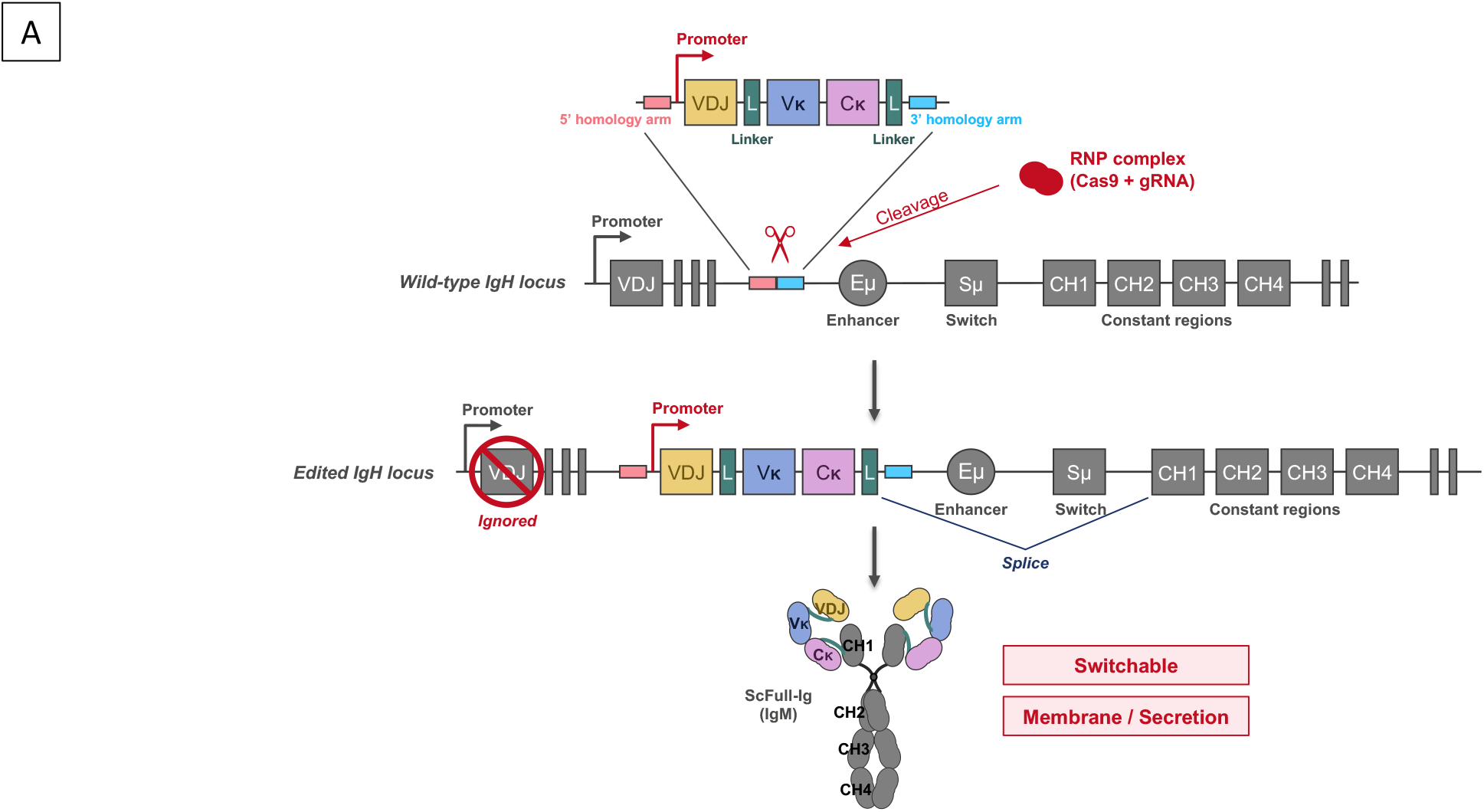
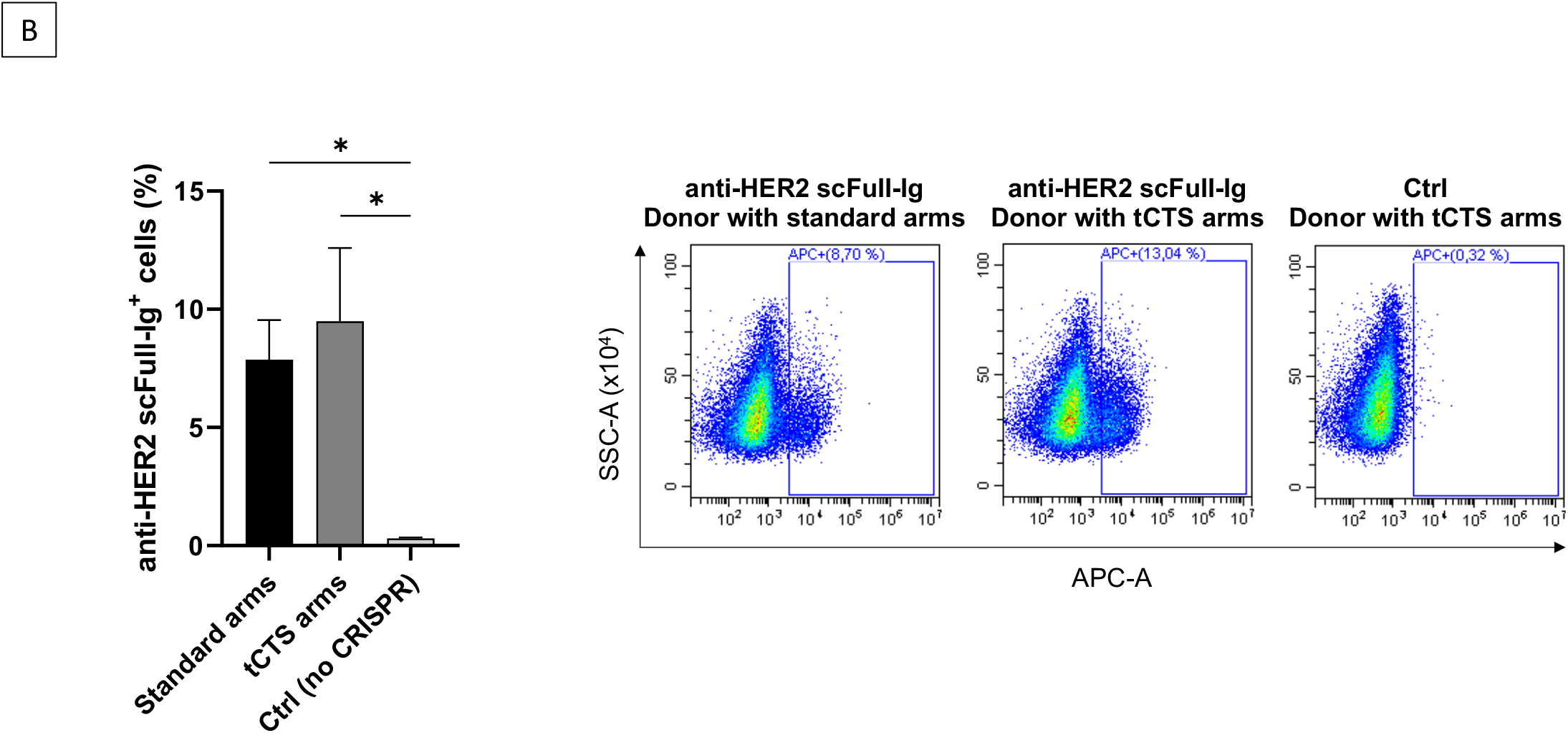
Knock-in in BL41 cell line. <u>(A)</u> Locus. The VDJ-L-Vκ-Cκ-L cassette is introduced between J_H_ and the Eμ enhancer with CRISPR-mediated cleavage. Insertion at this location using a V_H_ promoter allows expression of the edited Ig while disrupting expression of the endogenous Ig. Because the endogenous constant region is used, alternative splicing is not affected. The produced Ig can therefore be expressed as membrane-bound or secreted. All target regions for CSR are also preserved, while SHM could simultaneously affect the VH and Vk regions. <u>(B)</u> Expression of membrane-bound scFull-Ig. Cell surface staining for the anti-HER2 BCR was performed on gene edited BL41 cells, with His-tagged HER2 protein and APC-labelled anti-His antibody. tCTS, truncated Cas9 Target Sequence.

To further validate the strategy *in vitro*, we focused on the scFull-Ig targeting HER2, which displayed the highest functionality. We first tested the anti-HER2 scFull-Ig cassette in BL41 lymphoma cells with a mature B lymphocyte phenotype, featuring IgM isotype BCR expression. This experiment assessed the expression of the membrane-bound scFull-Ig IgM (Figure 2.B). After CRISPR-insertion of a [VDJ_pertu_-L-VκJκ_pertu_-Cκ-L-J_H_3’splice], around 8 to 9.5% of cells were successfully edited and acquired ability to bind soluble HER2, the best efficiency being again obtained, as for insertion of the tdTomato cassette, when HDR arms were ending with tCTS sequences promoting their binding (without cleavage) by the specific gRNA and Cas9.

To confirm the correct splicing of the HDR template on the endogenous IgM constant regions, we extracted the RNA of gene-edited cells at day seven from transfection and sequenced the IgH transcript after reverse transcription and amplification by PCR. One primer was designed to bind the scFull-Ig KI cassette and the other one the μ constant domain, allowing PCR amplification of the expected product only when the KI cassette correctly spliced with the endogenous constant domain. Correct splicing was observed, consistent with the membrane expression and functionality (Sup Figure 2.A).

Edition of BCR expression through the KI of an scFull-Ig cassette is thus efficiently promoted by CRISPR-mediated insertion in a human B cell line, prompting to extend this strategy to human primary B cells obtained from donors.

### Edition of Ig expression in primary B cells

Primary B cells were magnetically sorted from buffy coats and cultured for two days under our “1^st^ step stimulation cocktail” (including CpG, anti-BCR, CD40L and IL-2). At day two, CRISPR-mediated gene edition was carried out with the same conditions as for BL41 cells but with addition of Electroporation Enhancer and HDR Enhancer (IDT), following supplier’s instructions. Expression of anti-HER2 scFull-Ig at the cell membrane was evaluated at day seven. Around 3% of edited cells were then able to bind soluble HER2, the best efficiency obtained when using a donor DNA with tCTS sequences, with addition of poly-glutamic acid (PGA), Electroporation Enhancer and HDR Enhancer (Figure 3). As for BL41 cells, the correct splicing of the HDR template on the endogenous constant regions was confirmed by transcript analysis. At day seven, we confirmed correct RNA splicing of the scFull KI cassette onto endogenous constant Ig genes, corresponding to the expression of not only IgM but also class-switched isotypes such as IgA, as identified by RT-PCR with adequate specific primers (Sup Figure 2.B). Altogether, these data validate the fact that KI of the [VDJ_pertu_-L-VκJ_H_pertu-Cκ-L-J_H_3’splice] cassette in the IgH locus upstream of the Eμ enhancer allows expression of the scFull-Ig in primary B cells both in association with Cμ and with downstream class-switched isotypes.

**Figure 3.**
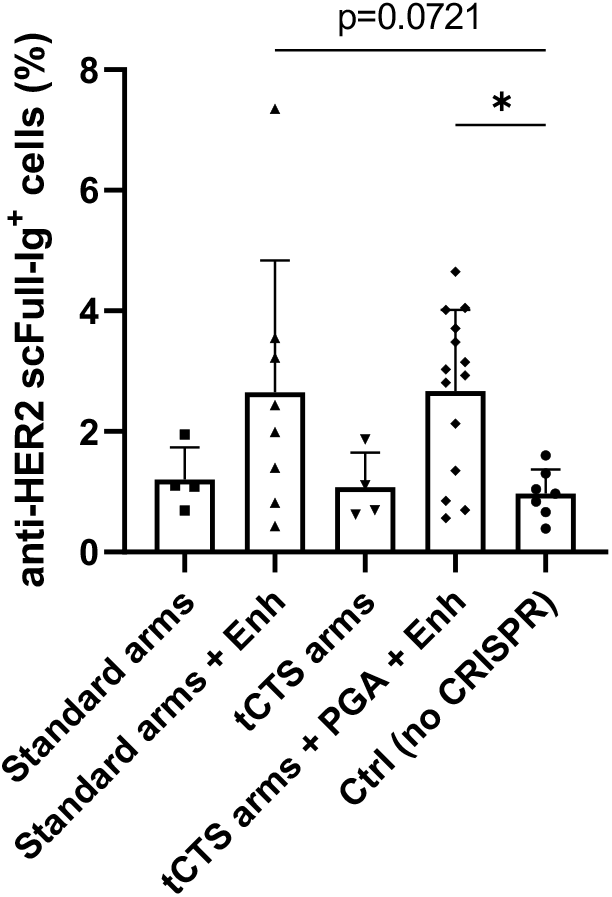
Knock-in in primary B cells, expression of the membrane-bound form. Cell surface staining for the anti-HER2 BCR was performed on edited human primary B cells, in the same way as for BL41 cells.

## DISCUSSION

Adoptive cell therapy has remarkably developed in the last decade. While T cells are currently the main actors of such strategies, B cells have recently received preliminary interest. B-lineage cells can abundantly secrete proteins and initial attempts to make them produce therapeutic proteins have relied on random lentiviral integration, notably for secreting Ig or coagulation factors^10,11^. For Ig, such strategies however expose to generating cells with dual specificity and/or with a mixture of desired and undesired H-L associations. CRISPR-edition of Ig loci is thus much more attractive since it can simultaneously terminate expression of the endogenous Ig and replace it with a mAb of interest^12–14^. In this regard, re-formatting the IgH locus is of special interest due to the competence of this locus for various spontaneous variations, with its notable ability to yield both membrane-bound and secreted Ig. Due to these intrinsic properties of the IgH locus, edited human B cells could thus fulfill two different and complementary therapeutic goals: either secreting a mAb of interest on the long term when differentiated into long-lived CD138^+^ CD20^-^ plasma cells, or patrolling in the patient’s lymphoid tissues as Ag-specific CD20^+^ CD27^+^ memory B cells to secrete the edited Ig only after Ag encounter. If done in a manner respecting the physiological organization of the human IgH locus, such an edition could even remain compatible with class-switching and then provide edited IgM molecules together with a variety of secondary class-switched variants carrying diversified constant regions, *i.e*. susceptible to exert a diversified spectrum of effector functions. Ultimately, if the edited V regions stand within the IgH locus territory accessible to SHM (*i.e*. within 1.5 kb downstream of V_H_ gene transcription initiation)^15^, the edited Ig could theoretically follow the path of affinity maturation and either increase its original affinity for the target Ag or get adapted to antigenic variations. The latter property would be of special interest for Ag showing spontaneous variations, as it occurs in chronic viral infections and in some cancer cells.

Previous studies have proposed single KI strategies in the IgH locus for expression of both the Ig H and L chains (eventually associating both chains covalently through a long linker peptide)^13,16^. However, they included some Ig constant sequences at an odd location belonging to the 1.5 kb upstream domain affected by SHM, immediately 3’ of the gene promoter^13,16^. By contrast, we elaborated a versatile strategy for also hijacking Ig production in B cells though a single IgH locus edition, based on the expression of scFull-Ig sequences starting with tandemly associated V_H_ and V_l_ sequences. In case edited B cells get activated, the scFull design should thus spare C exons from most of the SHM activity, and by contrast expose both the V_H_ and the V_l_ domains to affinity maturation. The scFull-Ig format proposed in this study thus combines several advantages. The single-chain design simplifies gene edition and warrants assembly of given IgH and IgL chains as a unique full Ig molecule. This format also complies with the objective of including both the V_H_ and V_l_ regions within an upstream transcribed IgH domain optimally exposed to high SHM in activated B lymphocytes. The other major expected outputs of this strategy, since based on IgH locus edition, are to allow expression of both membrane-associated BCR and secreted antibodies including an intact endogenous Fc domain, therefore expected to keep Ig effector functions unchanged.

This study aimed at validating functionality of such a single-chain format with regards to its capacity to be efficiently secreted, to maintain Ag specificity of the scFv domain while getting it finally associated with a classical and unmodified Fc domain. We indeed checked in the models of anti-CD20 rituximab and anti-HER2 pertuzumab, two classical therapeutic mAbs, that they kept the ability to efficiently induce ADCC against target cells when secreted under an scFull-IgG format.

Since insertion of linker elements within an Fv domain could eventually alter Ag binding, preservation of Ag specificity stands as an initial quality control for any mAb expressed in a single-chain format. We accordingly assessed preserved Ag binding in two models, based on heavy and light chain integration following the scFull design. Using CRIPSR-based edition of the IgH locus and KI of scFull-Ig cassettes, we confirmed in both models that both a human B cell line and human primary B cells can be efficiently gene modified for the expression of Ig: the upstream part of the cassette including both V regions and the C_l_ exon was correctly spliced onto the endogenous IgH C regions and edited B lymphocytes hereby expressed a BCR of the expected Ag specificity. Membrane-bound scFull-Ig efficiently interacted with the antigen and transcript sequencing revealed that scFull-Ig expressed in primary B cells was not only related to IgM, but also to class-switched isotypes such as IgA.

Many adjustments remain necessary before making edited human B cells good candidates for cellular therapy. One of the major issues is to enhance the efficacy of KI in this cell type. First of all, the KI efficiency relies on the delivery method of the DNA template to B cells, needing to combine high efficiency and low toxicity. Even if viral vectors such as Adeno-Associated Virus (AAV) are efficient carriers, we decided for safety reasons to focus our efforts on a simple virus free protocol, for which safe and well-designed production and purification methods should be easily transferable under Good Manufacturing Practices (GMP) conditions, as necessary for future preclinical development. Regarding safety, naked DNA-based strategies could indeed compete with other virus-free methods such as nanoparticle carriers^17,18^ or mAb encoded by mRNA^19^.

One limitation of B cell editing lies in the difficulty to perform HDR with exogenous DNA template at the optimal insertion site of the IgH locus. In order to optimize KI efficiency, we have checked the effect of varying the Cas9/sgRNA ratio as well as the amount of DNA template and we then sticked to the best conditions. Primary cell culture conditions prior and after the KI experiment were also worked out, in order to optimize both cell viability and HDR rates. Regarding the gene transfection itself, various HDR-enhancing molecules were tested in established cell lines and primary B cells, with the additional goal of limiting cell death after transfection. Based on Nguyen et al.^9^ publication, we were finally able to improve the rate of correct insertion at the IgH locus and the homogeneity of our results by using tCTS-modified dsDNA template, for routinely obtaining significantly high expression of the knock-in cassettes.

Altogether, while the protocol proposed in this study is quite efficient, further optimization will remain useful. In the future, strategies blocking the non-homologous end-joining (NHEJ) pathway^20^ might also deserve to be applied to targeted B cells in order to promote HDR. Both the HDR rate and the cell survival might also potentially be further improved by using ssDNA templates^21^, or by carrying pre-electroporation DNA-sensor inhibition, as reported for T cell editing^22^. Since B cells abundantly express intra-cellular Toll-like receptors, this could reduce the transfected DNA toxicity.

In conclusion, we report an efficient strategy to edit human B cells through a single precise gene modification of the IgH locus. This disrupts expression of the endogenous V_H_ domain and replaces it with a cassette providing most of the [Fv / C_L_] region from a mAb of interest, although in a single-chain format. Since this strategy rebuilds a complete Ig by reassociating the KI [Fv / C_L_] region to the endogenous [CH1 + Fc] part encoded by the IgH locus, it hereby warrants expression of the IgH chain under all possible normal variations, either expressed at the cell membrane as a BCR in lymphocytes or secreted by cells following plasma cell differentiation, or eventually switched of mutated once B cells have been activated. Edited B cells will then be ready for transfer into recipient mouse models, in order to assess antibody secretion and the potential adoptive humoral immunity.

## Notes

### Competing Interest Statement

The authors have declared no competing interest.

## References

1. Paci, A. et al. Pharmacokinetic/pharmacodynamic relationship of therapeutic monoclonal antibodies used in oncology: Part 1, monoclonal antibodies, antibody-drug conjugates and bispecific T-cell engagers. Eur J Cancer 128, 107–118 (2020).

2. Slifka, M. K., Antia, R., Whitmire, J. K. & Ahmed, R. Humoral Immunity Due to Long-Lived Plasma Cells. Immunity 8, 363–372 (1998).

3. Grupp, S. A. et al. Chimeric Antigen Receptor-Modified T Cells for Acute Lymphoid Leukemia. The New England Journal of Medicine 368, 1509–1518 (2013).

4. Jackson, H. J., Rafiq, S. & Brentjens, R. J. Driving CAR T-cells forward. Nat Rev Clin Oncol 13, 370–383 (2016).

5. Melenhorst, J. J. et al. Decade-long leukaemia remissions with persistence of CD4+ CAR T cells. Nature 602, 503–509 (2022).

6. Barrangou, R. & Doudna, J. A. Applications of CRISPR technologies in research and beyond. Nature Biotechnology 34, 933–941 (2016).

7. Johnson, M. J., Laoharawee, K., Lahr, W. S., Webber, B. R. & Moriarity, B. S. Engineering of Primary Human B cells with CRISPR/Cas9 Targeted Nuclease. Scientific Reports 8, (2018).

8. Le Gallou, S. et al. IL-2 Requirement for Human Plasma Cell Generation: Coupling Differentiation and Proliferation by Enhancing MAPK-ERK Signaling. J.I. 189, 161–173 (2012).

9. Nguyen, D. N. et al. Polymer-stabilized Cas9 nanoparticles and modified repair templates increase genome editing efficiency. Nature Biotechnology 38, 44–49 (2020).

10. Fusil, F. et al. A Lentiviral Vector Allowing Physiologically Regulated Membrane-anchored and Secreted Antibody Expression Depending on B-cell Maturation Status. Mol. Ther. 23, 1734–1747 (2015).

11. Levy, C. et al. Baboon envelope pseudotyped lentiviral vectors efficiently transduce human B cells and allow active factor IX B cell secretion in vivo in NOD/SCIDγc-/-mice. J. Thromb. Haemost. 14, 2478–2492 (2016).

12. Hartweger, H. et al. HIV-specific humoral immune responses by CRISPR/Cas9-edited B cells. J. Exp. Med. 216, 1301–1310 (2019).

13. Moffett, H. F. et al. B cells engineered to express pathogen-specific antibodies using CRISPR/Cas9 protect against infection. Science Immunol. 4, (2019).

14. Luo, B. et al. Engineering of α-PD-1 antibody-expressing long-lived plasma cells by CRISPR/Cas9-mediated targeted gene integration. Cell Death Dis 11, 973 (2020).

15. Peters, A. & Storb, U. Somatic hypermutation of immunoglobulin genes is linked to transcription initiation. Immunity 4, 57–65 (1996).

16. Hartweger, H. et al. HIV-specific humoral immune responses by CRISPR/Cas9-edited B cells. The Journal of experimental medicine 216, 1301–1310 (2019).

17. Nawaz, W. et al. Nanotechnology and immunoengineering: How nanotechnology can boost CAR-T therapy. Acta Biomater (2020) doi:10.1016/j.actbio.2020.04.015.

18. Smith, T. T. et al. In situ programming of leukaemia-specific T cells using synthetic DNA nanocarriers. Nat Nanotechnol 12, 813–820 (2017).

19. Pardi, N. et al. Administration of nucleoside-modified mRNA encoding broadly neutralizing antibody protects humanized mice from HIV-1 challenge. Nat Commun 8, 14630 (2017).

20. Fu, Y.-W. et al. Dynamics and competition of CRISPR-Cas9 ribonucleoproteins and AAV donor-mediated NHEJ, MMEJ and HDR editing. Nucleic Acids Res 49, 969–985 (2021).

21. Shy, B. R. et al. Hybrid ssDNA repair templates enable high yield genome engineering in primary cells for disease modeling and cell therapy manufacturing. 2021.09.02.458799 (2021) doi:10.1101/2021.09.02.458799.

22. Kath, J. et al. Pharmacological interventions enhance virus-free generation of TRAC-replaced CAR T cells. Mol Ther Methods Clin Dev 25, 311–330 (2022).

