## Supplemental information and figures for "Single-hit genome edition for expression of single-chain immunoglobulins by edited B cells"

### SUPPLEMENTARY INFORMATION

*BAR-B*

**Primer sequences**

| Primer | Sequence |
|---|---|
| Donor DNA amplification - For | TGGCCACTCTAGGGCCTTT |
| Donor DNA amplification - Rev | AGCTTGCTTTGGCCTCAATTC |
| Donor DNA amplification (tCTS) - For | TGGCGGGACTAGTGGCCCTTAGAGAACTGTCGGAGTGGGAA GAATGGCCACTCTAGGGC |
| Donor DNA amplification (tCTS) - Rev | TGGCGGGACTAGTGGCCCTTAGAGAACTGTCGGAGTGGGAG CTTGCTTTGGCCTCAATT |
| TIDE analysis - For | TGGGTTTTTGTGGGGTGAGG |
| TIDE analysis - Rev | GCTTTGGCCTCAATTCCAGAC |
| μ transcript analysis - RT | CGGGTRCTGCTGATGTCAGA |
| μ/α transcript analysis - PCR For | CCGCGGCCGCGCCACCATGGACTGGACCTG |
| μ transcript analysis - PCR Rev | CTCGTATCCGACGGGGAATT |
| α transcript analysis - RT | AAGTCCAGCACCACATAGGG |
| α transcript analysis - PCR Rev | GTCTCGTGGGCTCGGAGATGTGTATAAGAGACAGGCGAYGAC CACGTTCCCATCT |

**KI cassette sequences**

*Tomato*

AAGAATGGCCACTCTAGGGCCTTTGTTTTCTGCTACTGCCTGTGGGGTTTCCTGAGCATTGCAGGTTGGTCCTC
GGGGCATGTTCCGAGGGGACCTGGGCGGACTGGCCAGGAGGGGATGGGCACTGGGGTGCCTTGAGGATCTG
GGAGCCTCTGTGGATTTTCCGATGCCTTTGGAAAATGGGACTCAGGTTGGGTGCGTCTGATGGAGTAACTGAG
CCTGGGGGCTTGGGGAGCCACATTTGGACGAGATGCCTGAACAAACCAGGGGTCTTAGTGATGGCTGAGGAA
TGTGTCTCAGGAGCGGTGTCTGTAGGACTGCAAGATCGCTGCACAGCAGCGAATCGTGAAATATTTTCTTTAG
AATTATGAGGTGCGCTGTGTGTCAACCTGCATCTTAAATTCTTTATTGGCTGGAAAGAGAACTGTCGGACGGCC
GAAGCTTAAAAACCTCAGAGGATTTGTCATCTCTAGGCCTGCTCAGTAGAGGTTGCTATATAGCAGGGAAACA
TGCAAATAAGGCCTCTCTCTTCTCATGAAAACGAGTCCTGAACTAACCTTGAATCTGAAGCAAAGGGGATCAGC
CCGAGATTCTCATTCAGTGATCAACACTGAACACACATCCGCGGCCGCATGGTTTCCAAGGGTGAGGAGGTTA
TCAAAGAGTTCATGAGATTCAAGGTTAGGATGGAAGGTTCCATGAACGGTCACGAGTTCGAGATCGAGGGCG
AGGGTGAAGGTAGACCCTACGAGGGCACCCAAACCGCAAAGCTCAAAGTGACTAAGGGTGGTCCTTTGCCCT
TCGCTTGGGACATCTTGTCCCCCCAATTCATGTATGGCTCTAAGGCATACGTTAAGCATCCTGCTGACATCCCCG
ATTACAAAAAGTTGTCCTTCCCAGAGGGTTTCAAGTGGGAAAGGGTCATGAACTTCGAGGATGGAGGTCTTGT
GACTGTGACCCAAGATTCTAGTTTGCAGGACGGCACTTTGATCTACAAGGTGAAGATGAGAGGCACAAACTTT
CCTCCCGATGGTCCAGTCATGCAAAAGAAAACTATGGGTTGGGAAGCCTCCACTGAGAGGCTTTACCCAAGAG
ACGGCGTTCTTAAGGGTGAAATCCACCAAGCTCTCAAACTTAAGGATGGAGGCCACTACTTGGTGGAGTTCAA
GACCATCTACATGGCTAAGAAGCCCGTGCAACTCCCCGGCTATTACTACGTGGACACTAAACTCGATATCACCT
CCCACAACGAGGACTACACCATCGTTGAACAATATGAGAGGTCTGAGGGTCGCCATCACCTTTTCTTGGGTCAT
GGTACTGGAAGCACCGGTAGTGGCAGCTCTGGCACCGCTTCATCCGAGGATAATAACATGGCTGTGATCAAG
GAGTTTATGCGCTTCAAAGTCCGTATGGAGGGCTCAATGAATGGCCACGAGTTCGAGATCGAAGGAGAGGGT
GAGGGCCGCCCATATGAGGGCACTCAGACAGCTAAGTTGAAAGTCACCAAGGGTGGACCACTTCCTTTCGCTT
GGGATATTCTCTCACCACAGTTTATGTACGGTTCCAAGGCTTACGTGAAACACCCAGCCGATATTCCAGATTAT
AAGAAGTTGTCTTTCCCAGAAGGATTTAAGTGGGAGCGCGTTATGAACTTCGAGGACGGTGGTTTGGTTACAG
TCACCCAAGACTCCTCCCTTCAAGATGGTACGCTTATCTACAAGGTCAAAATGCGTGGAACCAATTTCCCACCA
GACGGCCCAGTTATGCAGAAGAAGACTATGGGCTGGGAGGCTTCAACAGAGCGCTTGTATCCCCGCGATGGA
GTGTTGAAGGGCGAGATTCACCAGGCATTGAAGTTGAAGGACGGTGGACATTACCTCGTGGAGTTTAAGACC

*BAR-B*

ATCTACATGGCCAAGAAACCCGTTCAATTGCCTGGTTATTACTACGTTGATACCAAGTTGGACATTACCTCCCAC
AACGAGGATTACACCATTGTCGAACAGTACGAGCGTTCCGAGGGCCGCCACCACCTCTTCCTCTACGGTATGG
ACGAGTTGTACAAGTGAAACTTGTTTATTGCAGCTTATAATGGTTACAAATAAAGCAATAGCATCACAAATTTC
ACAAATAAAGCATTTTTTTCACTGCATTCTAGTTGTGGTTTGTCCAAACTCATCAATGTATCTTAAGTGGGTGAA
TCCAGCCAGGAGGGACGCGTAGCCCCGGTCTTGATGAGAGCAGGGTTGGGGGCAGGGGTAGCCCAGAAACG
GTGGCTGCCGTCCTGACAGGGGCTTAGGGAGGCTCCAGGACCTCAGTGCCTTGAAGCTGGTTTCCATGAGAA
AAGGATTGTTTATCTTAGGAGGCATGCTTACTGTTAAAAGACAGGATATGTTTGAAGTGGCTTCTGAGAAAAA
TGGTTAAGAAAATTATGACTTAAAAATGTGAGAGATTTTCAAGTATATTAATTTTTTTAACTGTCCAAGTATTTG
AAATTCTTATCATTTGATTAACACCCATGAGTGATATGTGTCTGGAATTGAGGCCAAAGCAAGCT

*ScFull-Ig HER2*

AAGAATGGCCACTCTAGGGCCTTTGTTTTCTGCTACTGCCTGTGGGGTTTCCTGAGCATTGCAGGTTGGTCCTC
GGGGCATGTTCCGAGGGGACCTGGGCGGACTGGCCAGGAGGGGATGGGCACTGGGGTGCCTTGAGGATCTG
GGAGCCTCTGTGGATTTTCCGATGCCTTTGGAAAATGGGACTCAGGTTGGGTGCGTCTGATGGAGTAACTGAG
CCTGGGGGCTTGGGGAGCCACATTTGGACGAGATGCCTGAACAAACCAGGGGTCTTAGTGATGGCTGAGGAA
TGTGTCTCAGGAGCGGTGTCTGTAGGACTGCAAGATCGCTGCACAGCAGCGAATCGTGAAATATTTTCTTTAG
AATTATGAGGTGCGCTGTGTGTCAACCTGCATCTTAAATTCTTTATTGGCTGGAAAGAGAACTGTCGGACGGCC
GAAGCTTAAAAACCTCAGAGGATTTGTCATCTCTAGGCCTGCTCAGTAGAGGTTGCTATATAGCAGGGAAACA
TGCAAATAAGGCCTCTCTCTTCTCATGAAAACGAGTCCTGAACTAACCTTGAATCTGAAGCAAAGGGGATCAGC
CCGAGATTCTCATTCAGTGATCAACACTGAACACACATCCGCGGCCGCGCCACCATGGACTGGACCTGGAGCA
TCCTCTTCTTGGTGGCAGCAGCAACAGGTAAGGGGCTCCCCAGTCTCGGGGTTGAGGCAGAAACCAGGCCACT
CAAGTGAGGCTTTACCCACCCCTGTGTCCTCTCCACAGGTACCTACTCCGAAGTGCAGCTCGTCGAAAGCGGTG
GCGGACTGGTTCAGCCCGGTGGTTCTCTGCGGCTGTCTTGTGCTGCCTCGGGTTTCACGTTCACTGACTACACA
ATGGACTGGGTGCGTCAGGCTCCTGGAAAGGGATTGGAGTGGGTAGCCGACGTTAATCCAAACTCCGGCGGG
AGCATCTACAACCAGAGGTTCAAGGGGAGGTTCACTCTGAGCGTGGATCGCTCCAAGAACACGCTGTACCTCC
AGATGAACTCTCTCAGGGCCGAGGACACGGCTGTTTACTATTGCGCGAGGAACCTGGGTCCTTCCTTCTACTTC
GACTACTGGGGACAGGGAACCCTGGTGACCGTCAGCTCCGGTGGTGGTGGTTCTGGTGGTGGTGGTTCTGGC
GGCGGCGGCTCCAGTGGTGGTGGATCCGATATTCAGATGACCCAGTCCCCAAGCTCCCTGAGTGCCTCAGTGG
GCGACCGAGTCACCATCACATGCAAGGCTTCCCAGGATGTGTCTATTGGAGTCGCATGGTACCAGCAGAAGCC
AGGCAAAGCACCCAAGCTGCTGATCTATAGCGCCTCCTACCGGTATACCGGCGTGCCCTCTAGATTCTCTGGCA
GTGGGTCAGGAACAGACTTTACTCTGACCATCTCTAGTCTGCAGCCTGAGGATTTCGCTACCTACTATTGCCAG
CAGTACTATATCTACCCATATACCTTTGGCCAGGGGACAAAAGTGGAGATCAAGAGGACTGTGGCCGCTCCCT
CCGTCTTCATTTTTCCCCCTTCTGACGAACAGCTGAAAAGTGGCACAGCCAGCGTGGTCTGTCTGCTGAACAAT
TTCTACCCTCGCGAAGCCAAAGTGCAGTGGAAGGTCGATAACGCTCTGCAGAGCGGCAACAGCCAGGAGTCT
GTGACTGAACAGGACAGTAAAGATTCAACCTATAGCCTGTCAAGCACACTGACTCTGAGCAAGGCAGACTACG
AGAAGCACAAAGTGTATGCCTGCGAAGTCACACATCAGGGGCTGTCCTCTCCTGTGACTAAGAGCTTTAACAG
AGGAGAGTGTTCTGGTGGCGGTGGCTCGGGCGGAGGTGGGTCGGGTGGCGGCGGATCAGGTGAGTCCTCAC
AACCTCTCTCCTGCTTTAACTCTGAAGGGTTTTGCTGCATTTTTGGGGGGGGTGAATCCAGCCAGGAGGGACG
CGTAGCCCCGGTCTTGATGAGAGCAGGGTTGGGGGCAGGGGTAGCCCAGAAACGGTGGCTGCCGTCCTGAC
AGGGGCTTAGGGAGGCTCCAGGACCTCAGTGCCTTGAAGCTGGTTTCCATGAGAAAAGGATTGTTTATCTTAG
GAGGCATGCTTACTGTTAAAAGACAGGATATGTTTGAAGTGGCTTCTGAGAAAAATGGTTAAGAAAATTATGA
CTTAAAAATGTGAGAGATTTTCAAGTATATTAATTTTTTTAACTGTCCAAGTATTTGAAATTCTTATCATTTGATT
AACACCCATGAGTGATATGTGTCTGGAATTGAGGCCAAAGCAAGCT

**LEGENDS TO SUPPLEMENTAL FIGURES:**

**Supplemental Figure 1. tdTomato knock-in in BL41.**

A CRISPR-mediated tdTomato reporter gene KI was carried out in BL41 cells. With about 30% of tdTomato positive cells, we validated the possibility to insert a large expression cassette in this locus. The efficiency of tCTS-modified homology arms was also confirmed. tCTS, truncated Cas9 Target Sequence.

**Supplemental Figure 2. Validation of correct splicing of the knock-in exons on the endogenous IgH constant exons**

(A) In BL41 cells. RNA of edited cells was extracted, and the IgH transcript was sequenced after reverse transcription and PCR. Correct splicing of the inserted cassette on the endogenous constant region was confirmed.

(B) In primary B cells. RNA of edited primary B cells was extracted, and splicing was confirmed by performing RT-PCR on IgH transcripts, on unswitched Cμ gene or Cα gene after class-switching.

Sup Figure 1. tdTomato knock-in in BL41

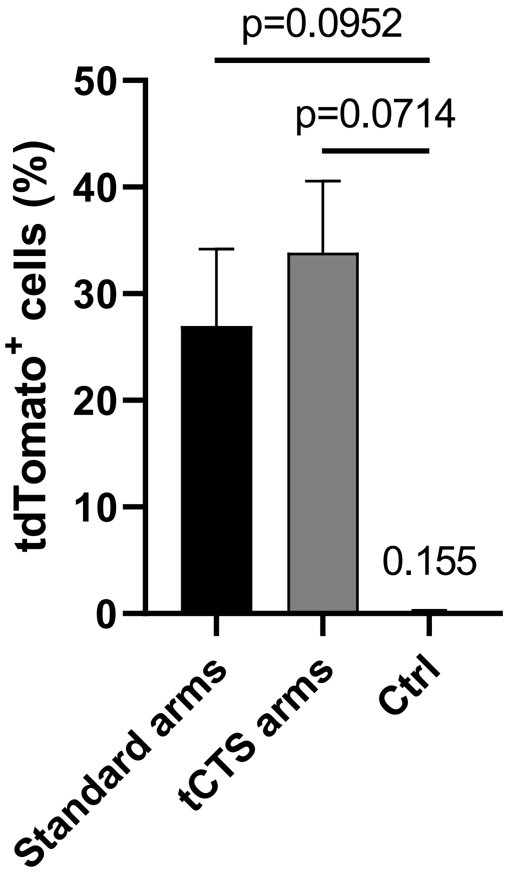

Sup. Figure 2. Validation of correct splicing of the knock-in exons on the endogenous IgH constant exons

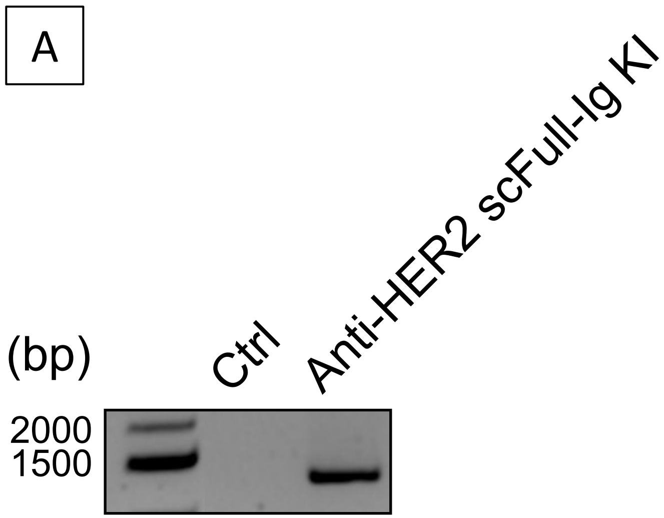


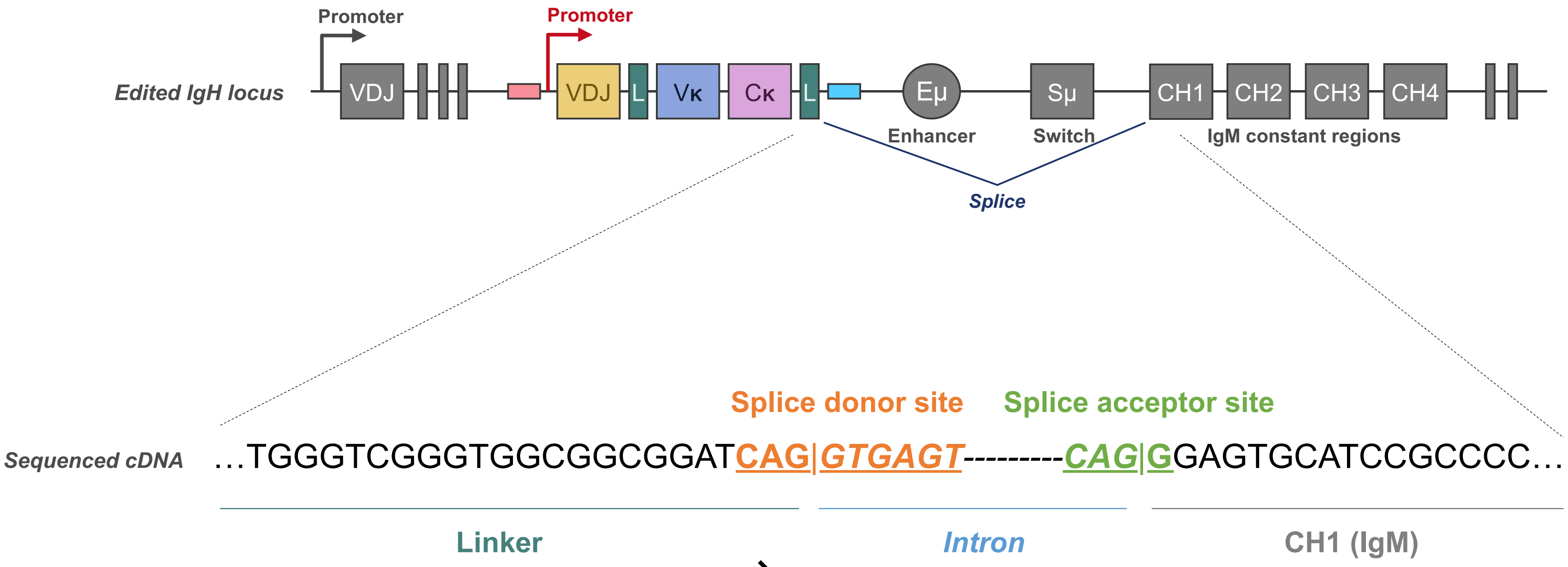


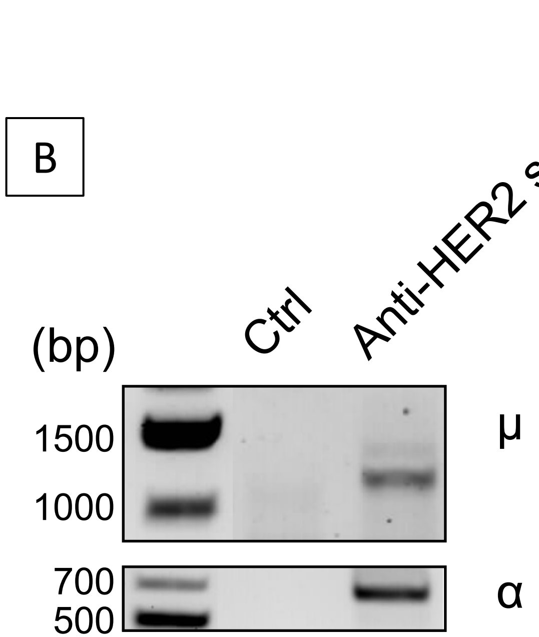
